# The structural evolution of host-pathogen protein interactions: an integrative approach

**DOI:** 10.1101/581637

**Authors:** Anderson F. Brito, John W. Pinney

## Abstract

The evolution of protein-protein interactions (PPIs) is directly influenced by the evolutionary histories of the genes and the species encoding the interacting proteins. When it comes to PPIs of host-pathogen systems, the complexity of their evolution is much higher, as two independent, but biologically associated entities, are involved. In this work, an integrative approach combining phylogenetics, tree reconciliations, ancestral sequence reconstructions, and homology modelling is proposed for studying the evolution of host-pathogen PPIs. As a case study, we analysed the evolution of interactions between herpesviral glycoproteins gD/gG and the cell membrane proteins nectins. By modelling the structures of more than 12,000 ancestral states of these virus-host complexes it was found that in early times of their evolution, these proteins were unable to interact, most probably due to electrostatic incompatibilities between their interfaces. After the event of gene duplication that gave rise to a paralog of gD (known as gG), both protein lineages evolved following distinct functional constraints, with most gD reaching high binding affinities towards nectins, while gG lost such ability, most probably due to a process of neofunctionalization. Based on their favourable interaction energies (negative ΔG), it is possible to hypothesize that apart from nectins 1 and 2, some alphaherpesviruses might also use nectins 3 and 4 as cell receptors. These findings show that the proposed integrative method is suitable for modelling the evolution of host-pathogen protein interactions, and useful for raising new hypotheses that broaden our understanding about the evolutionary history of PPIs, and their molecular functioning.

## INTRODUCTION

Despite being directly dependent on their hosts, the evolution of viruses does not always track the evolution of their hosts (de Vienne et al. 2013; Geoghegan et al. 2017). In a similar fashion, the evolutionary history of genes, although directly associated with their species evolution, may also follow their own independent pathways, such as duplications, losses and transfers (Page and Charleston 1998). In this scenario, the evolution of virus-host protein interactions is a systemic process governed by genetic and organismal phenomena.

To replicate, viruses establish several protein-protein interactions (PPIs) to hijack the cell machinery; and despite genetic differences, related viral species tend to use similar PPIs to accomplish this task (Brito and Pinney 2017). Subdivided in three subfamilies, *Alpha*-, *Beta*- and *Gammaherpesvirinae* (Davison et al. 2009), herpesviruses (HVs) are double-stranded DNA viruses with large genomes encoding dozens or even hundreds of genes (Brister et al. 2015). In alphaHVs, at least five genes are known to encode envelope glycoproteins involved in viral entry mechanisms, they are: gB, gC, gD, gH and gL (Spear and Longnecker 2003; Davison 2007). Among them, gD is responsible for binding cell receptors, and initiating a cascade of events leading to viral entry by fusion, or endocytosis followed by fusion (Krummenacher et al. 2005; Connolly et al. 2011). gD is only found in alphaHVs, and some species are known to encode a paralog of this gene, named gG (gene US4) (McGeoch et al. 1987), which diverged and developed the ability to block the activation of cell migration by functioning as a decoy receptor for chemokines (Bryant et al. 2003; Van de Walle et al. 2009). In cell entry mechanisms, nectins are the main cell receptors for alphaHVs. These immunoglobulin(Ig)-like proteins are expressed by various cell types, and act as transmembrane cell adhesion molecules (CAM) (Takai et al. 2008; Harrison et al. 2012). They are found in four types – nectin-1, nectin-2, nectin-3 and nectin-4 (Takai et al. 2008) –, and gD is known to interact with nectins 1 and 2, but is apparently unable to bind nectins 3 and 4 (Geraghty et al. 1998; Takai et al. 2008)

In light of the functional differences among such viral and host paralogs, by studying the evolution of their PPIs, new insights on the long-term evolutionary dynamics of herpesviruses and their hosts can be gained. Disagreements between viral and host phylogenies are common (Page and Charleston 1998; de Vienne et al. 2013), and for similar reasons, disagreements between gene trees and species trees are frequently observed, especially due to events of horizontal gene transfers, deletions and gene duplications (Page and Charleston 1998). By reconciling gene trees and time-calibrated species trees, the importance of such events can be assessed, and their times of occurrence can be assigned along the phylogenies of viral and host protein families (Conow et al. 2010). Another important approach for studying the evolution of PPIs is ancestral sequence reconstruction. Based on existing protein sequences, their gene trees, and substitution models, populations including the most likely ancestral protein sequences can be reconstructed for each internal node of the gene trees (Chang and Donoghue 2000). By combining this information with that provided by tree reconciliations, pairs of coexisting proteins can be directly associated to assemble ancestral protein complexes (Ashkenazy et al. 2012; Rouet et al. 2017). By using a template structure of a PPI containing homologs of those ancestral proteins, 3D structures of ancient PPIs can be obtained by homology modelling (Sali and Kuriyan 1999; Aloy et al. 2003). Importantly, despite low levels of sequence identity (30-40%), the structural properties of homologous proteins tend to be highly conserved, making ancestral sequence reconstruction and homology modelling useful resources for understanding the evolution of PPIs (Aloy et al. 2003; Rouet et al. 2017). By pairing ancestral virus-host homologs with distinct sequence compositions, the impact of mutations on their binding affinities can be directly assessed by using computational tools to measure structural properties such as interaction energy (Guerois et al. 2002) and electrostatic potentials (Jurrus et al. 2018).

Here, we propose a new method for studying the structural evolution of PPIs, which combines phylogenetics, tree reconciliations, ancestral sequence reconstructions, and homology modelling. As a study case, we applied this method to analyse the evolution of virus-host PPIs involving the herpesviral glycoproteins D/G and the cell adhesion proteins nectins. Our analyses revealed how gD and gG probably evolved their distinct molecular functions, and based on measures of binding affinity, we hypothesize that apart from nectins 1 and 2, other nectins may also be involved in cell entry mechanisms of alphaHVs. Overall, our methodology proved to be robust, being not just applicable for analysing evolving host-pathogen protein pairs, but also other kinds of PPIs.

## METHODS

### Viral and host species and their proteins

As gD (domain ‘Herpes_glycop_D’) is exclusively found in *Alphaherpesvirinae*, only species from this subfamily and their respective host species were included in this study (Table 1). Sequences of orthologs and paralogs of the protein pair found in the template structure PDB 3U82 (Zhang et al. 2011) were retrieved from NCBI. To identify gD and its paralog (glycoprotein G), searches for the domain ‘Herpes_glycop_D’ were performed against translated ORFs from all fully sequenced HV genomes using HMMER-3.1 (Mistry et al. 2013). Likewise, four sets of proteins from the Nectin family (Nectin-1, −2, −3, and −4) were retrieved via Blast searches (Altschul et al. 1990), using as query the human Nectin-1 found in the template structure (Figure 1A) (Zhang et al. 2011). Only one copy of each Nectin type was included per host species, and when no sequence was available, sequences from the closest available species were used.

**Table 1.** Alphaherpesviruses and hosts included in this study.

| Genus | Virus | Viral species | Host species | Host |
| --- | --- | --- | --- | --- |
| *Iltovirus* | GaHV1 | *Gallid alphaherpesvirus 1* | *Gallus gallus* | Gga |
| - | PsHV1 | *Psittacid alphaherpesvirus 1* | *Amazona oratrix* | Psi |
| *Mardivirus* | AnHV1 | *Anatid alphaherpesvirus 1* | *Anser cygnoides* | Acy |
| - | CoHV1 | *Columbid alphaherpesvirus 1* | *Columba livia* | Cli |
| - | FaHV1 | *Falconid herpesvirus 1* | *Falco mexicanus* | Fal |
| - | GaHV2 | *Gallid alphaherpesvirus 2* | *Gallus gallus* | Gga |
| - | GaHV3 | *Gallid alphaherpesvirus 3* | *Gallus gallus* | Gga |
| - | MeHV1 | *Meleagrid alphaherpesvirus 1* | *Meleagris gallopavo* | Mga |
| - | SpHV2 | *Spheniscid herpesvirus 2* | *Spheniscus* | Sph |
| *Scutavirus* | TeHV3 | *Testudinid herpesvirus 3* | *Testudo horsfieldii* | Tho |
| *Simplexvirus* | CeHV1 | *Macacine alphaherpesvirus 1* | *Macaca mulatta* | Mma |
| - | CeHV16 | *Papiine alphaherpesvirus 2* | *Papio cynocephalus* | Pap |
| - | ChHV1 | *Chimpanzee herpesvirus strain 105640* | *Pan troglodytes* | Ptr |
| - | FBaHV1 | *Fruit bat alphaherpesvirus 1* | *Pteropus sp.* | Pte |
| - | HHV1 | *Human alphaherpesvirus 1* | *Homo sapiens* | Hsa |
| - | HHV2 | *Human alphaherpesvirus 2* | *Homo sapiens* | Hsa |
| - | LHV4 | *Leporid alphaherpesvirus 4* | *Oryctolagus cuniculus* | Ocu |
| - | SaHV1 | *Saimiriine alphaherpesvirus 1* | *Saimiri sp.* | Sai |
| *Varicellovirus* | BoHV1 | *Bovine alphaherpesvirus 1* | *Bos taurus* | Bta |
| - | BoHV5 | *Bovine alphaherpesvirus 5* | *Bos taurus* | Bta |
| - | CaHV1 | *Canid alphaherpesvirus 1* | *Canis lupus familiaris* | Cfa |
| - | CeHV9 | *Cercopithecine alphaherpesvirus 9* | *Erythrocebus patas* | Chl |
| - | EHV1 | *Equid alphaherpesvirus 1* | *Equus caballus* | Eca |
| - | EHV3 | *Equid alphaherpesvirus 3* | *Equus caballus* | Eca |
| - | EHV4 | *Equid alphaherpesvirus 4* | *Equus caballus* | Eca |
| - | EHV8 | *Equid alphaherpesvirus 8* | *Equus caballus* | Eca |
| - | EHV9 | *Equid alphaherpesvirus 9* | *Equus caballus* | Eca |
| - | FeHV1 | *Felid alphaherpesvirus 1* | *Felis catus* | Fca |
| - | HHV3 | *Human alphaherpesvirus 3* | *Homo sapiens* | Hsa |
| - | SuHV1 | *Suid alphaherpesvirus 1* | *Sus scrofa* | Ssr |

**Figure 1.**
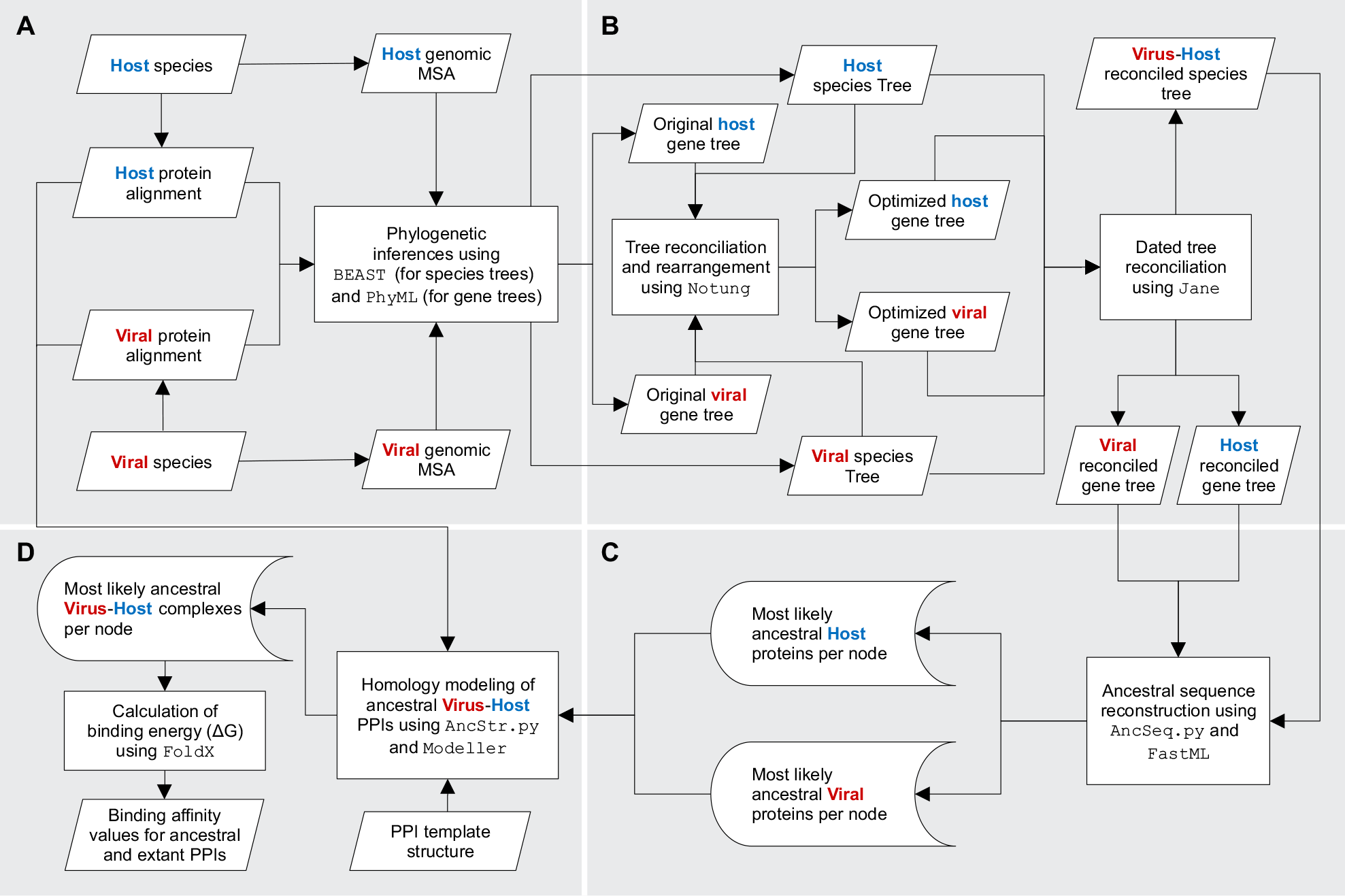
Pipeline for reconstruction of ancestral virus-host protein-protein interactions. A) Sequences and Phylogenies: once the host and viral species are selected, multiple sequence alignments (MSAs) of interacting proteins are generated for inferring their gene trees using PhyML. Similarly, informative genomic sequences are aligned for inferring viral and host species trees using BEAST. B) Tree reconciliations: in a species-to-species tree reconciliation, the viral species tree is reconciled with the host species tree using Jane. Next, two gene-to-species tree reconciliations are performed, with viral and host gene trees being reconciled in two steps, one for rearranging weak branches following the species tree topology, using NOTUNG, followed by another one, now between the optimized gene trees and their respective time-calibrated species tree, using Jane. C) Ancestral sequences: using information from the reconciliations, the optimized gene trees, and the protein alignments described in (A), the most likely ancestral sequences are reconstructed from each internal node of both gene trees. For this step, the software FastML is used to generate ungapped sequences, and a python script is used to assign gaps following probabilities per site provided as a FastML output. D) Ancestral PPIs: using the ancestral sequences reconstructed in (C) and an existing template structure, homology modelling is applied for reconstructing ancestral and present-day virus-host complexes. Finally, the binding affinities of these structural protein interactions are calculated using FoldX.

### Phylogenetic analyses

Each protein dataset was converted into multiple sequence alignments (MSAs) using Promals3D (Pei and Grishin 2007), which allowed us to generate protein alignments guided by structural information of the template complex (Figure 1A) (Zhang et al. 2011). The best fitting amino acid substitution models were determined using Protest (Darriba et al. 2011), and the evolutionary relationships of host and viral proteins were inferred using PhyML (Guindon et al. 2010). To understand the evolution of host and viral species alongside their respective interacting protein families, time-calibrated species phylogenies were inferred using a Markov Chain Monte Carlo (MCMC) Bayesian approach implemented in BEAST v2.4.5 (Bouckaert et al. 2014). The host tree was inferred using nucleotide MSAs of the nuclear genes BDNF, CNR1, EDG1, RAG1, and RHO, with their substitution models being determined using jModelTest (Posada 2008). The viral tree was reconstructed using amino acid MSAs of the core proteins encoded by UL15, UL27 and UL30, with substitution models defined by ProtTest (Darriba et al. 2011). These alignments were used as partitions in *BEAST. Priors for node ages were calibrated using divergence times according to (Hedges and Kumar 2009; dos Reis et al. 2012; Claramunt and Cracraft 2015), for host species; and according to (McGeoch and Gatherer 2005; Wertheim et al. 2014), for viral species. The host tree was run for 500 million generations, and the viral one for 35 million generations, both under relaxed (uncorrelated lognormal) molecular clocks, with Yule model as coalescent prior.

### Tree reconciliations

Despite being biologically associated, mismatches between the phylogenies of host and viral species are common. For similar reasons, the same principle applies to the phylogenetic history of proteins and organisms, as genes also undergo duplications, transfers and losses (Page and Charleston 1998). To unravel the role of such events in the evolution of the species and their interacting proteins, two types of tree reconciliations were performed: species-to-species; and gene-to-species reconciliation (Figure 1B). Firstly, the viral tree was reconciled with the host tree using the software Jane. Secondly, each protein/gene tree was reconciled with their respective species tree in two steps. The step 1 was carried out using the software Notung (Chen et al. 2000), which allows low support branches (bootstrap < 0.5) in gene trees to be rearranged to match its species tree topology, providing more parsimonious inferences of gene duplications and horizontal gene transfers. These improved gene trees served as input in the step 2 of reconciliation, using the software Jane (Conow et al. 2010). In this process, events of interspecific coalescence, gene duplications and losses were inferred and mapped along each species phylogenies. In both steps of reconciliations, the event costs were defined as follows: codivergences = 0; duplications = 1; transfers = 3; and losses = 1.

### Ancestral sequence reconstruction

Using the gene trees with improved topologies, the amino acid substitution models, and the datasets of present-day protein sequences, we performed ancestral sequence reconstructions using the software FastML (Figure 1C) (Ashkenazy et al. 2012). As the topology of gene trees were rearranged in the first step of reconciliation, their branch lengths were recalculated by FastML using the same substitution models applied for inferring their initial trees. Using a Perl script provided by its developers, the 10 most likely ancestral sequences at each internal node were generated. As these sequences are reconstructed without indels, a python script was developed to add gaps to such sequences following their probability of occurrence per site, according to the FastML output for marginal reconstruction of indels.

### Homology modelling

For homology modelling of ancestral PPIs, the results of tree reconciliations and ancestral sequence reconstructions were integrated. With the species-to-species reconciliation, ancestors of HVs were assigned to their most likely ancestral host lineages. As the repertoires of nectins and glycoproteins D/G in these ancestral virus-host pairs varied along their evolution, their gene-to-species reconciliations revealed how many copies of such proteins these species encoded in early times. Taking into consideration this chronological information, ancestral Nectin-gD/G complexes were modelled by homology using as template the structure PDB 3U82 (Zhang et al. 2011), and the structural protein alignments generated in Promals3D (Pei and Grishin 2007). All possible coexisting virus-host ancestral proteins were modelled in a pairwise manner using a python script calling MODELLER 9.18 (Figure 1D) (Sali and Blundell 1993). From five replicates generated per sequence pair, we selected the ones showing the lowest Discrete Optimized Protein Energy (DOPE score), which is calculated using an atomic distance-dependent statistical potential generated based on known native protein structures (Shen and Sali 2006). As the 10 most likely ancestral sequences were reconstructed per internal node in both protein trees, a total of 100 ancestral protein pairs were modelled per virus-host node association. Thereby, more than 63,000 ancestral complexes were modelled, with the top 12,500+ being selected for interaction energy calculations. In a similar way, present-day protein pairs (egg. SuHV1-gD/Suid-Nectin1) were also modelled, generating 165 complexes homologous to the template complex.

### Interaction energy and electrostatic calculations

Before calculating their interaction energy, each protein complex underwent energy minimization using FoldX/RepairPDB, which fixes residues with bad torsion angles and clashes, and finds their energy minimum at equilibrium. With the repaired virus-host complexes, using FoldX (Guerois et al. 2002; Schymkowitz et al. 2005) their interaction energy (ΔG_*bind*_, in kcal/mol) was calculated as follows:

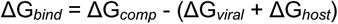

where ΔG_*comp*_ is the global free energy of the complex upon binding, ΔG_*viral*_ is the individual free energy of unfolding of the viral protein, and ΔG_*host*_ corresponds to such measure for the host protein. After Poisson-Boltzmann electrostatics calculations performed by PDB2PQR (Dolinsky et al. 2007), the electrostatic potentials of protein surfaces were obtained using the AMBER field force implemented on APBS (Figure 1D) (Jurrus et al. 2018).

## RESULTS

### Virus-host species tree reconciliation

To understand how virus-host protein-protein interactions evolve, it is essential to determine what viral lineages infected ancestral host species, which can be achieved by reconciling host and viral species trees. In this analysis, cospeciation is identified when a viral lineage co-diverges alongside their host species. When a deeper viral divergence is observed along the branches of the host tree, an intrahost speciation can be suggested. Using taxonomic information from host species, Figure 2 highlights how species tree reconciliation points out historical virus-host relationships. This information was crucial for establishing chronologically consistent ancestral protein pairs, as shown in the following sections.

**Figure 2.**
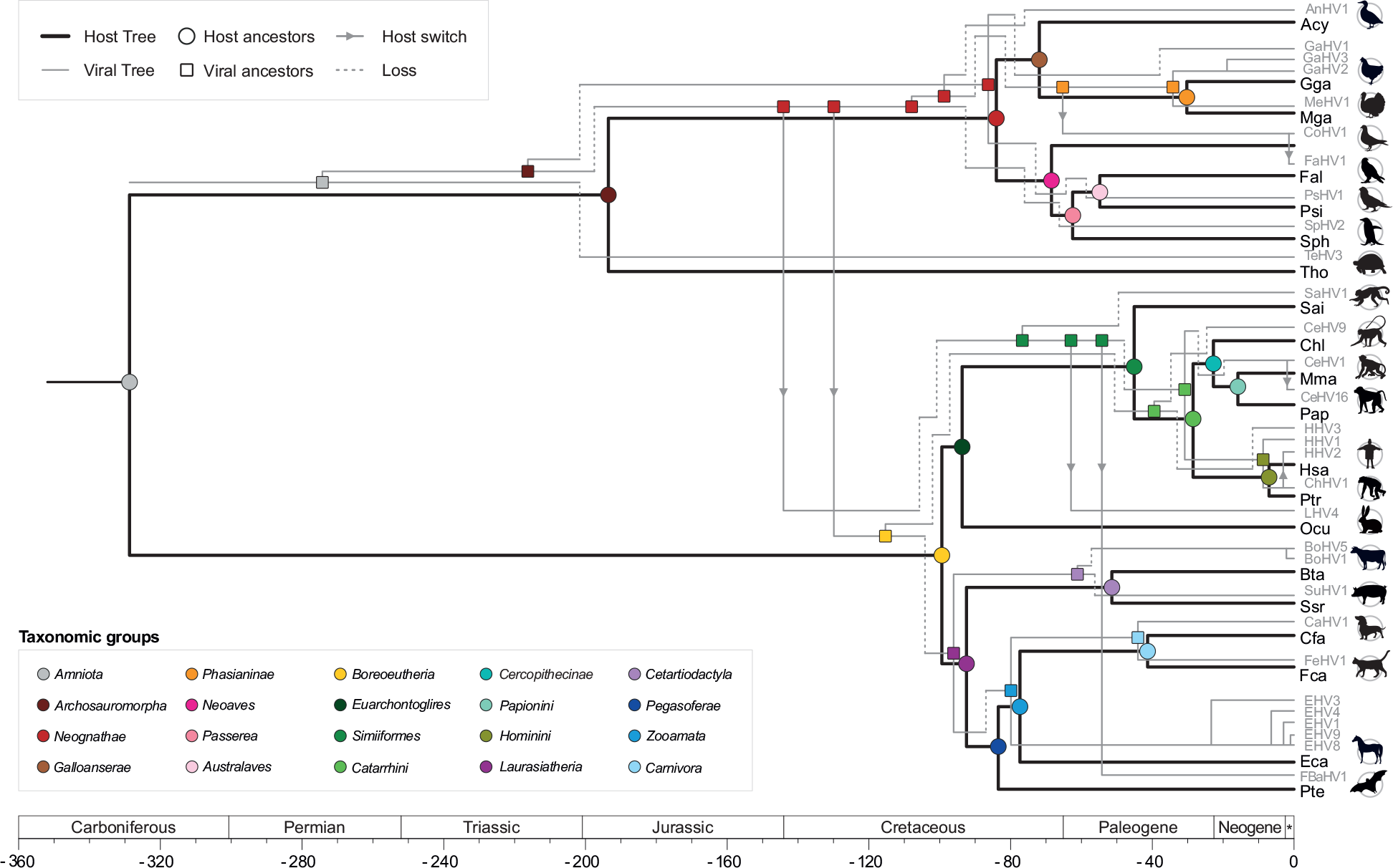
Virus-host species tree reconciliation. By reconciling viral and host species trees, internal nodes (i.e. ancestral states) of alphaHVs could be assigned along their host phylogeny, highlighting events of cospeciation, host transfers and intrahost speciations. Each node in the host tree is colour-coded according to their taxonomic classification, and those coloured with the same colours denote ancestral virus-host relationships.

### Host species and nectin gene tree reconciliation

Since events of gene duplication, transfers and deletions can generate gene tree topologies that disagree with those of their containing species (Page and Charleston 1998), to study the evolution of PPIs, reconciliations between gene and species trees should be performed to determine how genes evolved alongside species. To do so, gene trees with reliable topologies and correct rooting are essential. To root the host gene tree, non-nectin proteins encoding V-set domains were used as outgroup, which positioned the nectin-3 clade as the most basal group in this gene family (Figure S1A).

However, giving the fact that some branches had low support (bootstrap < 50%), before performing the final dated reconciliation using Jane, those weak branches were rearranged in a reconciliation using Notung, which uses the species tree topology as a guide to provide alternate and more robust hypotheses for its gene tree branching (Chen et al. 2000). By doing so one not just avoid the overestimation of duplication and loss events caused by inaccurate topologies, but also give more reliability to potential horizontal gene transfers (HGTs). In the gene tree of the nectin family, a total of 28 low support branches were rearranged after the initial reconciliation using Notung (Figure S1B). In this process, the tree root was repositioned splitting nectins in two groups: Nectins 1 and 2, and Nectins 3 and 4. After using this optimized tree for a final dated reconciliation using Jane (Figure 3), we identified that duplications of nectins were especially observed in the Carboniferous period (before ~350 Mya). Ancestors of mammals (*Boreoeutheria*) and birds (*Neognathae*) likely encoded several paralogs of nectins in that period (Figure 3), but many of them were either lost along the evolution or were not sampled in this study, since only a single copy of each nectin type was included, and many host genomes are not fully sequenced. Concerning gene losses, interestingly, our analysis shown that Nectin-2 was probably lost in *Archosauromorpha* ancestors in the Early Jurassic, after the split between ancestors of modern birds and reptiles (Figure 3). Finally, as expected for animal species, no HGTs of nectin genes were observed.

**Figure 3.**
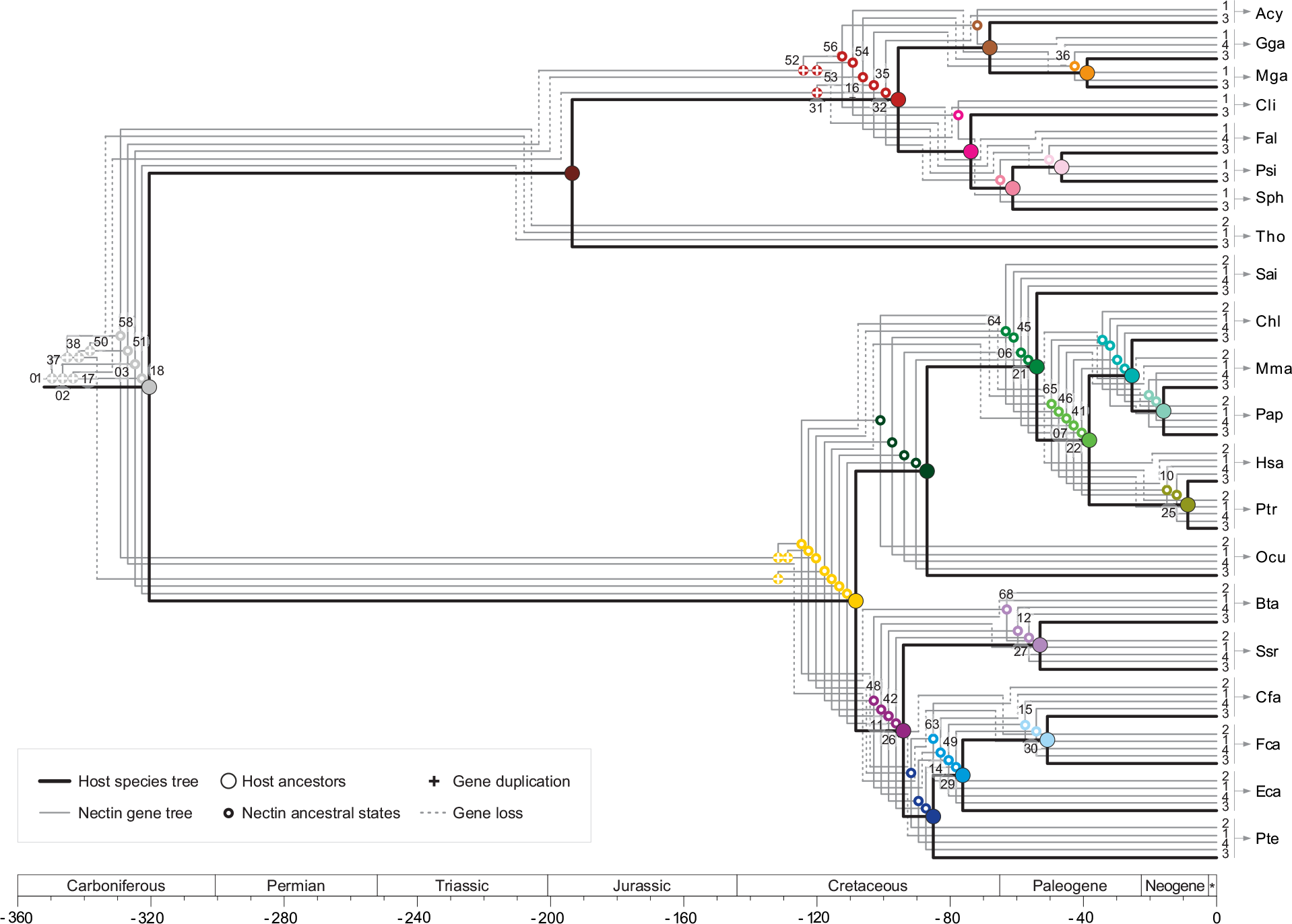
Reconciliation between the optimized nectin tree, and its species tree. Events of gene interspecific coalescence (gene-species divergence), gene duplication (deep coalescence) and gene losses (deletions or sorting events) are highlighted following the same colour-coding of Figure 2. For clarity, some internal nodes in the reconciled nectin tree are numbered.

### Viral species and glycoprotein D/G tree reconciliation

The gD/gG gene tree has shown better overall support than nectin tree, with most branch showing bootstrap values higher than 70%, and only 9 branches showing support below 50% (Figure S2A). As these viral genes are only encoded by *Alphaherpesvirinae*, no outgroup was available to resolve the polarity of these characters, and their tree was initially mid-rooted, splitting gD and gG in two monophyletic clades. After the first step of reconciliation for tree rearrangement, the repositioning of weak branches revealed a less costly hypothesis of tree rooting, which defined the gD of TeHV3 as the most basal taxon (Figure S2B). In this way, it minimized the number of events (losses and duplications) necessary to explain gD/gG evolution. This optimized gene tree was used in a final dated reconciliation, which showed that the last common ancestor of alphaHVs encoded a single copy of glycoprotein D, revealing only two duplications in this gene family (Figure 4). In the Triassic period, the first duplication gave rise to glycoprotein G, a paralog that was subsequently lost in *Iltovirus*, and is now only observed in *Mardivirus*, *Simplexvirus* and *Varicellovirus*. The second duplication took place most likely in the Early Cretaceous, where ancestors of alphaHVs infecting avian hosts gained another copy of gG, which was subsequently lost in ancestors of *Varicellovirus*, and deleted in multiple independent events among members of *Mardivirus* (Figure 4). Since only alphaHVs with fully sequenced genomes were considered in this study, these events of gene loss can be directly interpreted as gene deletions, making varicelloviruses infecting primates (CeHV9 and HHV3) the only species that lost all copies of gD/gG, most likely in the Early Cretaceous (~120 Mya). Finally, concerning HGTs, no transfers of gD/gG were found.

**Figure 4.**
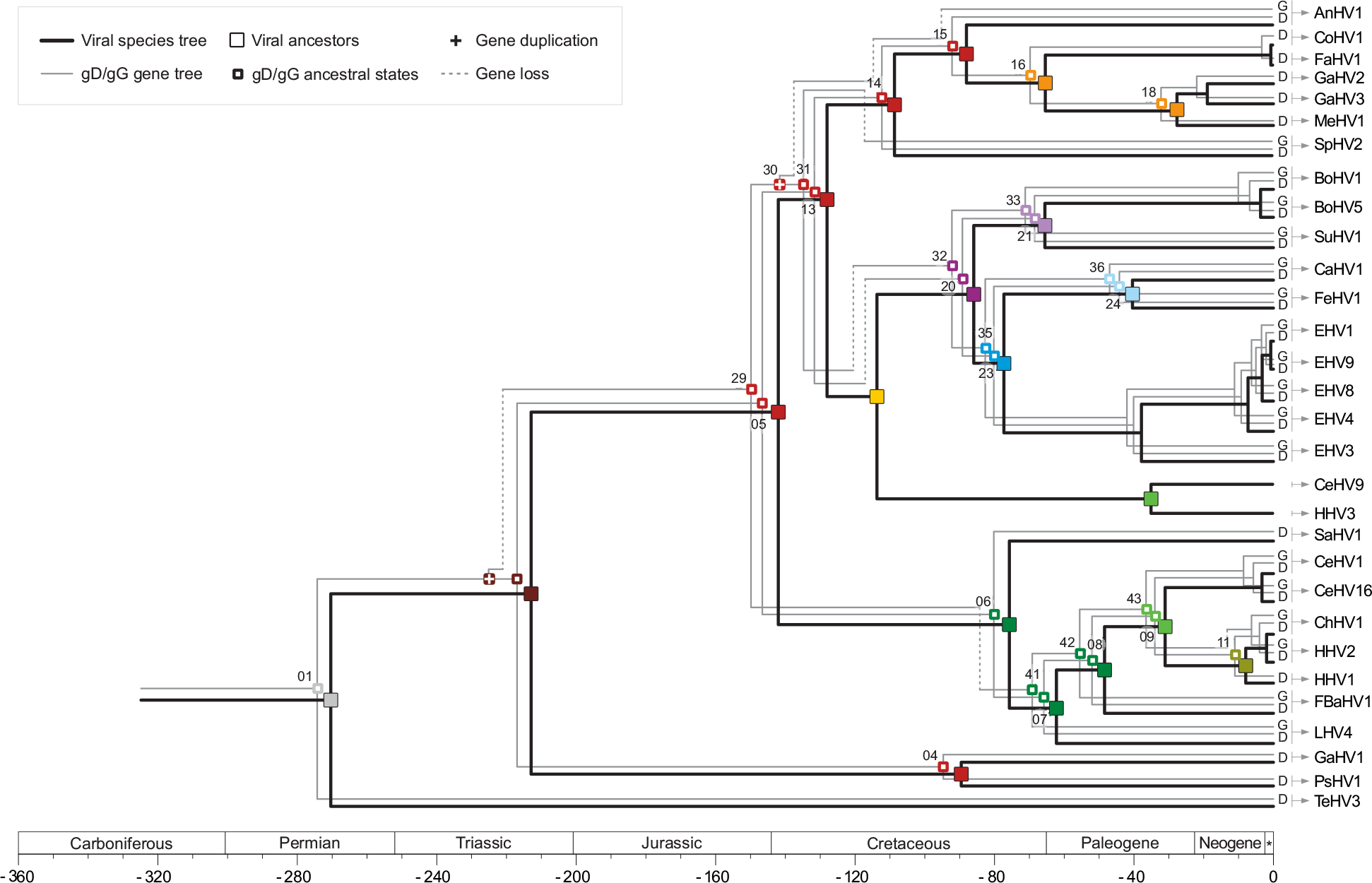
Reconciliation between the optimized gD/gG gene tree, and its species tree.

### Associating clusters of ancestral protein pairs

The reconciliation between gene and species tree determined the number of copies of nectins and glycoproteins D/G encoded by ancestral virus-host pairs. Based on this information, clusters of coexisting proteins were associated as ancestral virus-host PPIs, allowing us to generate structural models of these complexes by homology, and trace their evolution (Figure 5). From both gene trees, only internal nodes identified as interspecific coalescences were considered for homology modelling, because their divergence times coincide with their species divergences. Thereby, nodes representing gene duplications were excluded from the homology modelling because their divergence times could not be directly inferred. The only exception to this rule was applied for the MRCA of all complexes. As only sequences at internal nodes could be reconstructed, after the reconciliation between gene and species tree, many clusters of ancestral host proteins found no corresponding partner in the viral side, and vice-versa, preventing us from reconstructing all possible ancestral PPIs. However, despite this limitation, many other ancestral complexes could be modelled for further structural comparisons.

**Figure 5.**
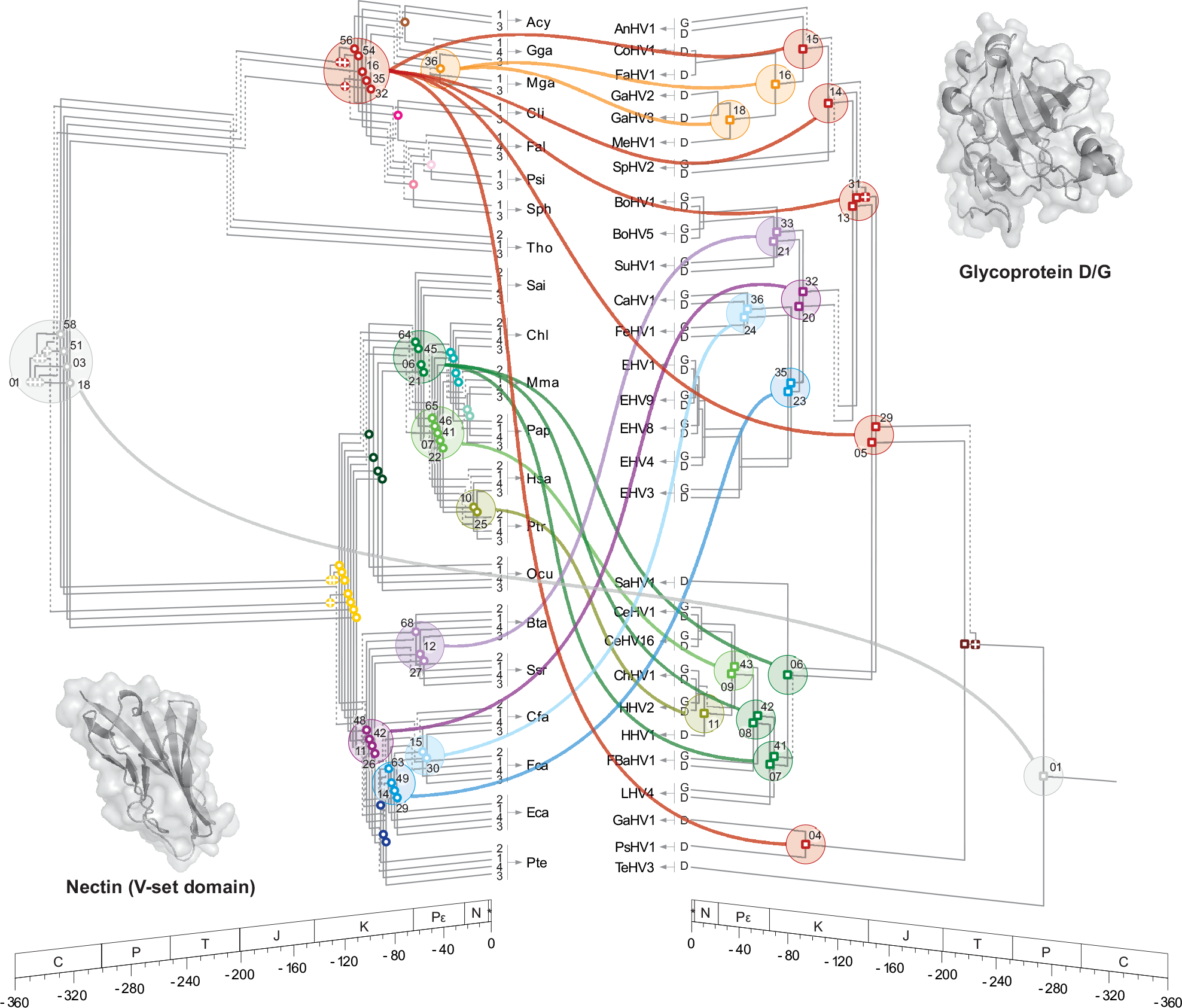
Clusters of ancestral protein pairs. In this representation, the reconciled gene trees found in Figure 3 and Figure 4 are shown in perspective. A total of 10 ancestral sequences were reconstructed for each of the numbered internal nodes. Sequences belonging to the same colour-coded clusters were pairwise associated for homology modelling of ancestral complexes, yielding 100 structures for each binary node association.

### Interaction tree: the evolution of PPIs

From the pairwise node associations shown in Figure 5, interaction trees depicting distinct evolving protein pairs were created as proposed by (Pinney et al. 2007) (Figure 6). To investigate how these PPIs evolved, their interaction energies (ΔG_*bind*_) were used as a proxy to capture changes on protein-protein affinity. To calculate the ΔG_*bind*_, Fold-X solves an energy function that includes several terms, such as: van der Waals contributions; hydrogen bonds; solvation energy for polar and polar groups; electrostatic contribution of charged groups, among others (Guerois et al. 2002). In energetic terms, a PPI is thermodynamically favourable if the ΔG of the complex (ΔG_*comp*_) is lower than the sum of the ΔG of the unbound forms of the interacting proteins. In this way, the lower the ΔG_*bind*_ of a protein pair, the higher the affinity between the interaction partners, with negative values representing processes that release energy (exergonic), and positive values representing processes that require absorption of external energy (endergonic) (Nelson and Cox 2012). In this way, protein pairs showing positive ΔG_*bind*_ probably are unable to interact.

**Figure 6.**
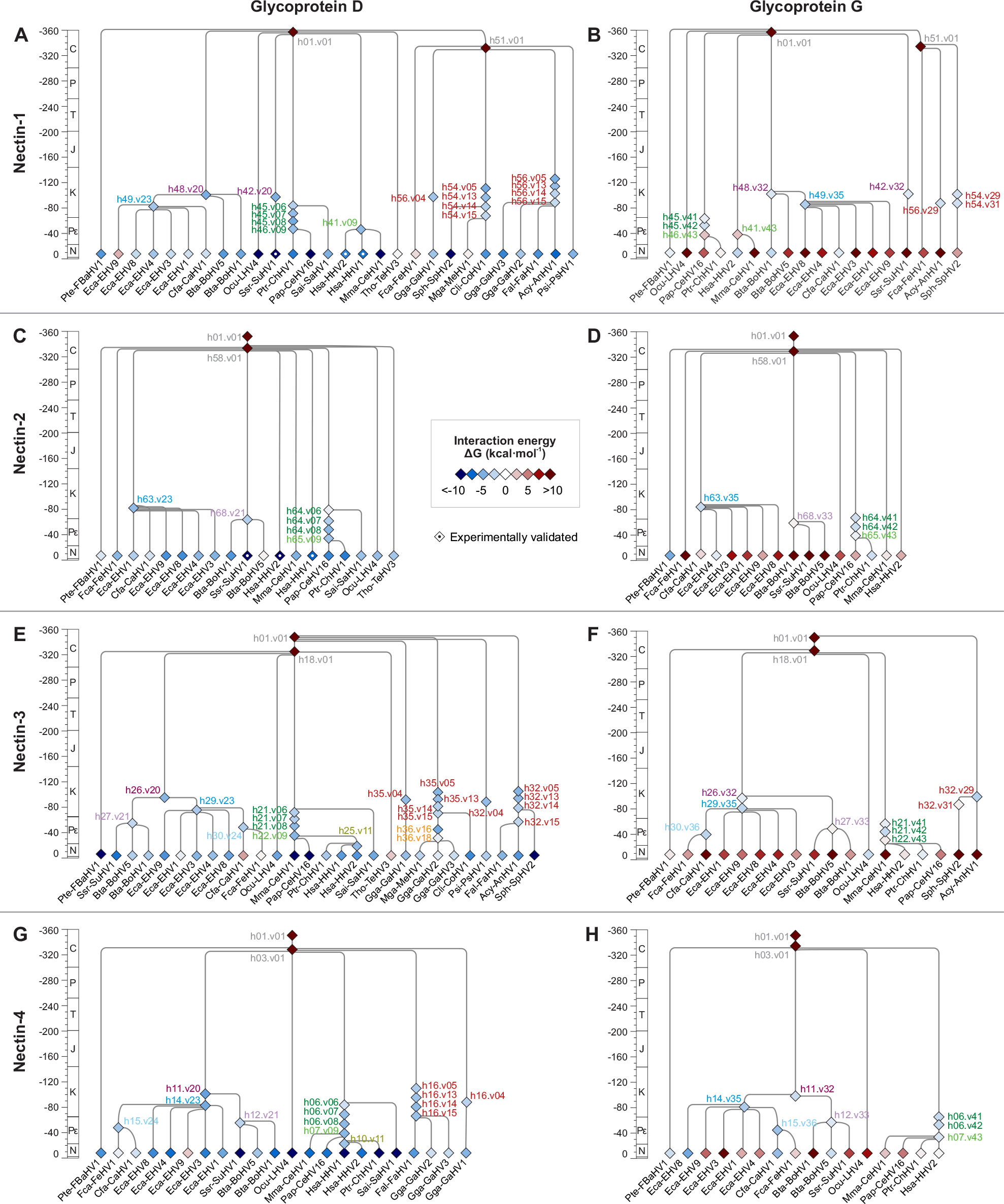
Interaction trees illustrating the evolution of virus-host PPIs. These trees show how the interaction energy (ΔG_*bind*_) of ancestral pairs of Nectins and gD/gG evolved, with each diamond-shaped node representing a PPI. The interaction energies shown at internal nodes come from the average energy of 100 structures, as explained in Figure 5. PPIs at the tips represent present-day protein-pairs, and their interaction energies come from a single replicate. PPIs that were experimentally validated are highlighted with an internal circle (•). A) Interaction tree of Nectin-1 and glycoprotein D. B) Nectin-1 and glycoprotein G. C) Nectin-2 and gD. D) Nectin-2 and gG. E) Nectin-3 and gD. F) Nectin-3 and gG. G) Nectin-4 and gD. H) Interaction tree of Nectin-4 and gG. Labels at internal nodes are colour-coded as in Figure 2, and their numbers correspond to those shown in Figure 5, i.e., ‘h48.v20’ represents an interaction between the viral gD (node 20) and the host nectin-1 (node 48) found in *Laurasiatheria* ancestors.

Starting from the last common ancestors of these PPIs, the complexes h01.v01, our results revealed that in early times these protein pairs had very low affinity. It showed an average ΔG_*bind*_ of 33.88 (± 32.62) kcal/mol, with 89% of the replicates showing positive ΔG_*bind*_ values, indicating an energetic incompatibility between these ancestral protein pairs (Figure 6). By looking at the electrostatic potential of the interfaces of these ancient proteins (Figure 7), which date from the Carboniferous period (C, ~310 Mya), it is possible to observe that the MRCA of gD/gG had largely positively charged interfaces, a property that could cause repulsions between these proteins and ancestral nectins. As most events of gene interspecific coalescence revealed by our study date from more recent times (Figure 5), it was not possible to trace exactly how the binding affinities and electrostatic potentials of these complexes evolved from the Devonian (D) to the Cretaceous (K) period. However, at least from the Cretaceous period to present times, it is possible to observe that both gD and gG gradually changed their ΔG_*bind*_ in opposite directions, respectively enhancing and weakening their affinities with nectins (Figure 7). This becomes clearer when the average ΔG_*bind*_ of intermediate states of present-day gD and gG are compared. The binding affinities of gD with nectins were, on average, higher (−3.75 < ΔG_*bind*_ < −3.00) than those involving gG (−2.09 < ΔG_*bind*_ < −0.76), differences that correlate well with the binding affinity of present-day complexes. The ΔG_*bind*_ of glycoproteins D towards all nectin types have shown negative values, indicating energetically favourable interactions (Figure 6A, C, E, and G). The average ΔG_*bind*_ of existing complexes between gD and nectin-1 and −2 were −5.52 and −5.45, respectively; while for nectin-3 and −4 the average ΔG_*bind*_ were −4.73 and −7.00 kcal/mol, respectively. In this way, the gD affinity towards nectin-4 was higher than those observed for nectins 1 and 2.

**Figure 7.**
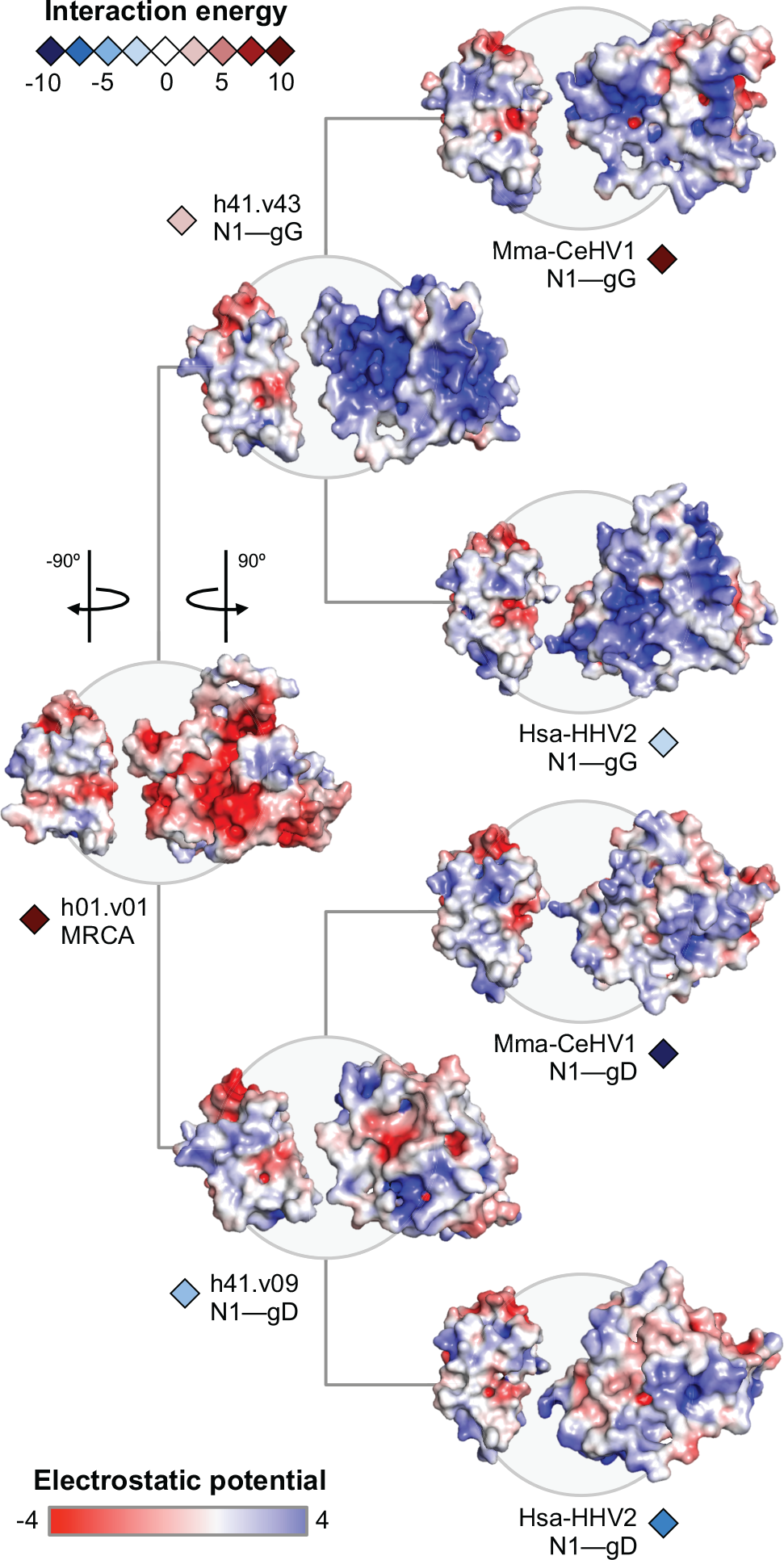
The evolution of interface electrostatic potentials. These complexes belong to HVs infecting Humans (HHV2) and Macaques (CeHV1), also represented in Figure 6A and B. In this illustration, ancestral states and existing PPIs are shown with their structures rotated 90º so that their interfaces are exposed. The interfaces of the gD/gG MRCA were mostly negatively charged, fact that may explain their low affinity (high ΔG_*bind*_) towards nectins. These host proteins, on the other hand, kept their interfaces with similar electrostatic patches along the evolution. Glycoproteins G diverged developing positively charged interfaces, while glycoproteins D evolved interfaces with more balanced distribution of charges, resembling those of nectins, which may explain their higher binding affinity (low ΔG_*bind*_).

Concerning the extant gG-nectin complexes, as expected, the opposite trend was observed, with most complexes showing neutral or positive ΔG_*bind*_ values (Figure 6B, D, F, H and Figure 8). Among the present-day gG complexes, the average binding energy varied from 3.86 (for gG-nectin4), to 18.10 kcal/mol (for gG-nectin3), revealing a clear energetic incompatibility between these protein pairs (Figure 6B, D, F, H). Most likely after the first duplication that gave rise to ancestors of glycoproteins G, most of these proteins decreased their binding affinity with nectins, probably due to changes in the electrostatic potential of their interfaces, which became largely positive by the Cretaceous (~110 Mya) (Figure 7). These findings match what is known about glycoproteins G, which probably lost their ability of cell receptor binding in a process of neofunctionalization (Bryant et al. 2003).

**Figure 8.**
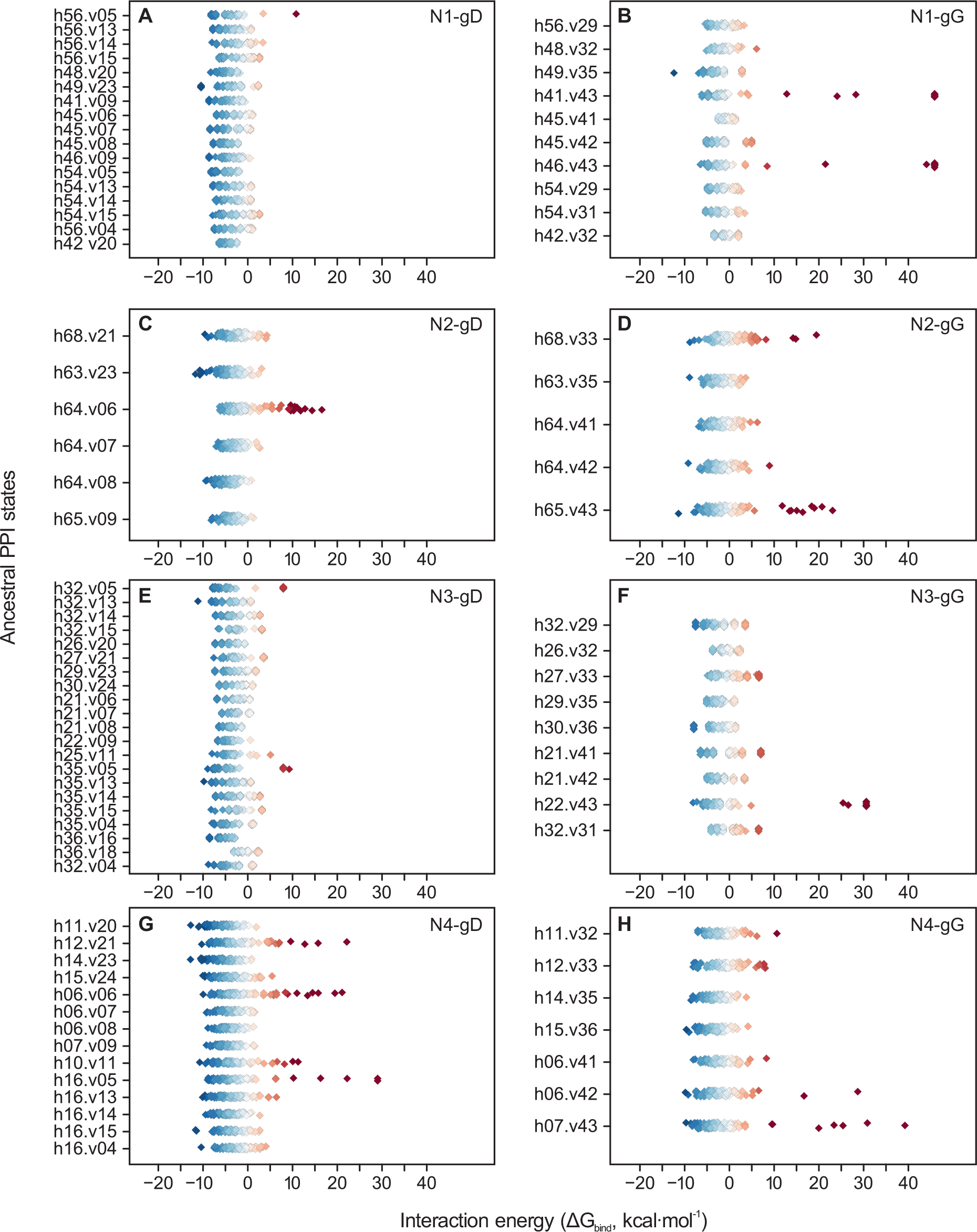
Distribution of binding affinities of ancestral complexes. For each internal of the interaction tree shown in Figure 6 a total of 100 replicates had their interaction energy calculated. It is possible to see a clear difference between gD complexes –with ΔG_*bind*_ fluctuating more towards negative (energetically favourable) values and gG complexes which show more neutral and positive (energetically unfavourable) values.

## DISCUSSION

In this study we present the results of an integrative approach for studying the evolution of host-pathogen PPIs. To develop and apply this method, homology modelling was applied to generate 3D structures of complexes obtained via ancestral sequence reconstruction. For such, we used as template a structurally solved interaction between glycoprotein D and Nectin1, which is found in HHV1-Human systems (PDB 3U82) (Zhang et al. 2011). Existing homologs of these interacting proteins were extracted from several virus-host pairs, and by reconciling their gene trees and respective time-calibrated species trees, it was possible to understand how these individual protein families evolved. More importantly, we could also identify ancestral protein pairs that probably coexisted and potentially interacted in the past, aspect that allowed us to reconstruct their ancestral protein-protein interactions. A total of 10 ancestral protein variants were reconstructed for each internal node of both gene trees, and their pairwise association as protein complexes allowed us to generate 100 PPI structures per ancestral state, resulting in over 12,000 homology models. This extensive analysis revealed that in the past, sequence variants of gD and gG had different binding affinities with nectins (Figure 8), and under distinct constrains, their evolutions were characterized by gradual changes on interaction energy between intermediate ancestral states.

By using measures of interaction energy calculated using FoldX (Guerois et al. 2002), our method was able to capture fine differences of binding affinity in complexes involving nectins and two lineages of viral glycoproteins. In early times of their evolution, such viral proteins were mostly unable to interact with nectins, probably due to electrostatic incompatibilities (Figure 7). Glycoproteins D/G encode a single domain ‘Herpes_glycop_D (PF01537)’, which is a member of the Immunoglobulin superfamily (CL0011) (Cocchi et al. 1998; Finn et al. 2014). Based on our co-phylogenetic analysis, this domain was acquired by alphaherpesviruses between the Permian and Carboniferous periods, and although its origin is still unclear, in light of its structural similarities with immunoglobulins, a hypothesis of acquisition via HGT from host cells is a plausible explanation. Since newly acquired proteins can take millions of years to adapt and integrate into an existing protein interaction network (Lercher and Pal 2008), as soon as gD was acquired, it was probably maladapted to establish interactions with nectins. However, as it evolved and duplicated, gD and its paralog (gG) optimized their affinities with their respective interaction partners following distinct evolutionary paths.

After the duplication event that gave rise to gG, this paralog probably had the ability to weakly bind nectins, however, most of them probably lost such affinity in more recent times (Figure 6B, D, F, H). Since duplicated genes may evolve under distinct functional constraints and substitution patterns (Zhang 2003), gG probably lost their ability to bind nectins due to their neofunctionalization as chemokine binding proteins (Bryant et al. 2003). In fact, these changes in binding affinity were probably important for maintaining both paralogs in the protein repertoires of alphaHVs, inasmuch as paralogs are unlikely to be kept in genomes if they perform identical functions (Hughes 1994; Zhang 2003). This aspect can for example explain the deletion of gG in some HVs, as observed in HVs infecting birds (genera *Iltovirus* and *Mardivirus*), and in the simplexviruses SaHV1 and HHV1 (Figure 4). Notably, among members of the genus *Varicellovirus*, HHV3 (Varicella-Zoster Virus) and CeHV9 (Simian Varicella Virus) are the only two species that do not encode gD and gG (Davison and Scott 1986; Gray et al. 2001)most probably due to deletions that took place along the Early Cretaceous. The reasons for such gene losses are still unclear, however, the deletion of gD highlights the high level of plasticity of these viruses, which circumvented the essentially of nectin-gD interactions (Kinchington et al. 2012) by means of alternative cell entry mechanisms exploring other cell receptors (Ouwendijk and Verjans 2015).

Nectins play a central role in the formation of cell-cell junctions (Takai et al. 2008). To perform their function, they establish self-interactions, and mutations at their interfaces can lead to severe anomalies (Rikitake et al. 2012), contexts in which purifying selection may act to purge deleterious mutations, preserving the compatibility between nectin interaction interfaces (Daugherty and Malik 2012). Based on the structure of HHV1-gD/nectin-1 complex, it was found that gD competes with other nectins by binding the same interface used by them to establish interactions (Zhang et al. 2011). In a context of evolutionary arms race, mutations at nectin interfaces would not just affect their self-interactions, but also the ability of gD to use them as cell receptors. At a population level, and in response to constant genetic conflicts, the binding affinities between these proteins may have evolved by gradual increases or decreases on interaction energy, with ancestral gD-nectin complexes fluctuating at much higher affinities (negative ΔG_*bind*_) than ancestral complexes of gG (Figure 6 and Figure 8). This observation correlates well with the ΔG_*bind*_ of present-day complexes, which despite being measured using a single replicate per complex, have shown similar overall ΔG_*bind*_ values (Figure 6).

Supported by experimental data, and showing 96% of accuracy, a previous computational model of binding affinity revealed that protein pairs showing ΔG_*bind*_ lower than −7 kcal/mol are very likely to form stable complexes (Kiel et al. 2005). Moreover, it also showed that protein pairs with ΔG_*bind*_ > −5 kcal/mol are probably unable to establish physiologically stable interactions (Kiel et al. 2005). In our analysis, by looking at the transitions between ancestral states and present-day protein pairs, it was possible to identity many instances where proteins expected to interact evolved towards much higher ΔG_*bind*_ (> 0 kcal/mol), aspect that may indicate loss of interaction in current virus-host associations. This trend is observed, for example, for the N1-gD pairs Eca-EHV9, Fca-FeHV1, and Mga-MeHV1 (Figure 6A), and for the N2-gD pair Bta-BoHV5 (Figure 6C), which probably are unable to establish interactions. Unfortunately, we could not find specific results validating the absence of these interactions. However, previous studies have validated the occurrence of some interactions, which match our findings based on ΔG_*bind*_, they are: the N1-gD interactions of Hsa-HHV1 (Zhang et al. 2011), Hsa-HHV2 (Lu et al. 2014), Ssr-SHV1 (Li et al. 2017); and the N2-gD interactions of Hsa-HHV1, Hsa-HHV2, and Ssr-SHV1 (Warner et al. 1998). Figure 6A and C highlight these experimentally validated PPIs with internal circles (•).

Examples of interaction loss were also observed for N3-gD and N4-gD, however, for these proteins the scenarios of interaction gain were more frequent, with many protein pairs evolving towards ΔG_*bind*_ below the threshold of −7 kcal/mol, indicating increases of binding affinity over time. Such high affinities were particularly observed in the N3-gD complexes Pte-FBaHV1; Mma-CeHV1; Pap-CeHV16; Sph-SpHV2 (Figure 6E); and in the N4-gD complexes Ssr-SuHV1; Ocu-LHV4; Hsa-HHV1; Ptr-ChHV1; and Sai-SaHV1 (Figure 6G, Figure 8G). By examining the binding energies of gD and different nectin types, we found results that disagree with prior assumptions. Despite no experimental analysis have directly investigated the binding affinities of Nectins 3 and 4 with glycoproteins D, manystudies have incorrectly implied that there is evidence against these proteins acting as cell entry receptors for herpesviruses (Struyf et al. 2002; Krummenacher et al. 2003; Petermann et al. 2015), while other studies proposed further analysis to confirm or reject such hypothesis (Reymond et al. 2000; Satoh-Horikawa et al. 2000). Our interaction energy models revealed that some gD may interact with nectin-3, and more importantly, with nectin-4, as this receptor showed the highest overall affinity with gD. Nectin-4 is known to be involved in cell entry mechanisms of paramyxoviruses (Mühlebach et al. 2011; Noyce et al. 2013; Delpeut et al. 2014; Singh et al. 2016), but so far, no study reported its use by herpesviruses. The aforementioned N4-gD protein pairs, as well as those involving Nectin-3, are good candidates for experimental validation, and further researches investigating these interactions would be highly beneficial, especially if they include virus-host pairs other than the commonly studied.

Despite the current low availability of viral protein structures, as the number of 3D structures of virus-host complexes grows, evolutionary analysis of additional PPIs could shed light on how other fundamental interactions of viral infection cycles evolved. From a systemic perspective of protein interaction networks, being able to predict a large number of PPIs would ultimately allow us to assess, for example, the chances a new viral strain has to complete its infection cycle, taking into consideration its overall protein binding affinities with host proteins involved in several intermediate steps of the infection, from cell entry to virion egress. By integrating phylogenetics, ancestral sequence reconstruction, tree reconciliations and structural biology, in this study were generated more than 12,000 PPIs between coexisting ancestral proteins from distinct virus-host pairs to study their evolution. This method proved to be suitable for this purpose, being applicable for evolutionary studies of any protein-protein interaction, especially for those of host-pathogen systems. To evaluate the level of feasibility of this method, an interaction involving members of two diverse gene families was used as a study model: gD/gG and nectins. The lower binding affinities of gG with nectins were in accordance with previous studies, since these glycoproteins gained new functions along their evolution. However, unexpected results were found concerning the binding affinities of gD with nectins 3 and 4, which were previously suggested not to be herpesvirus receptors. As similar results were found across distinct virus-host pairs, including their ancestral states, these findings indicate that more thorough investigations on such protein interactions would be extremely advantageous for validating or rejecting our hypothesis of new receptors for alphaherpesviruses.

## ACKNOWLEDGMENTS

AFB is funded by *Ciência sem Fronteiras*, a scholarship programme managed by the Brazilian federal government (CAPES, Ministry of Education, Grant number: 11911-13-1). JWP is supported by a University Research Fellowship from the Royal Society.

## Authors contributions

AFP and JWP conceived and designed the study; analyzed the data and wrote the manuscript.

---

**Figure S1.**
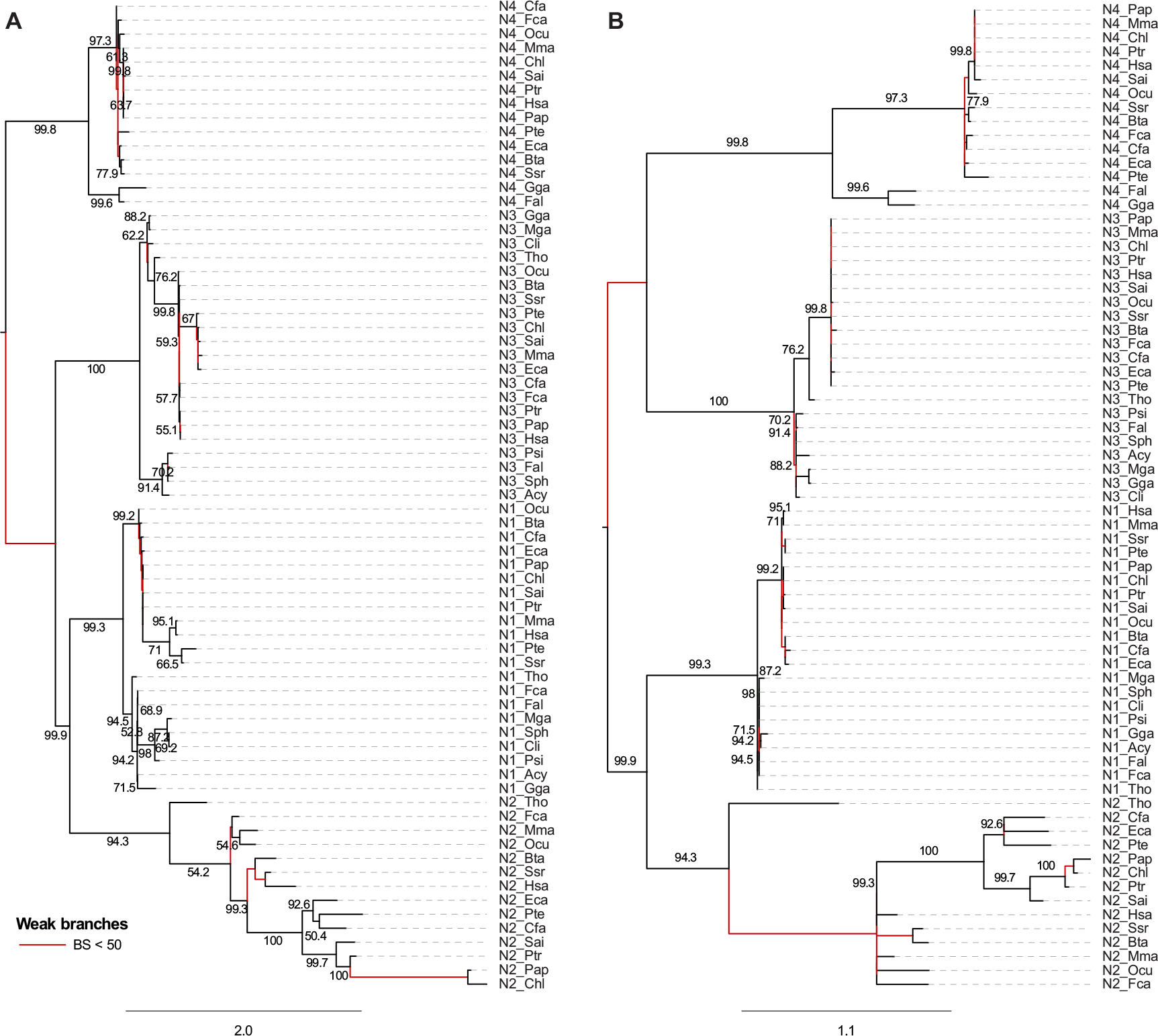
Nectin gene trees. A) Original tree inferred using PhyML. Two membrane proteins homologous to nectins were used as outgroup (Uniprot E1BMI5 and Q549Q4, not shown in the trees). Branches with bootstrap support below 50% are coloured in red. These weak branches were rearranged to match its host species tree topology in a first step of reconciliation using Notung. B) Optimized nectin gene tree, with the topology of strongly supported branches kept as in the original tree shown in (A), and new branching pattern for weak branches.

**Figure S2.**
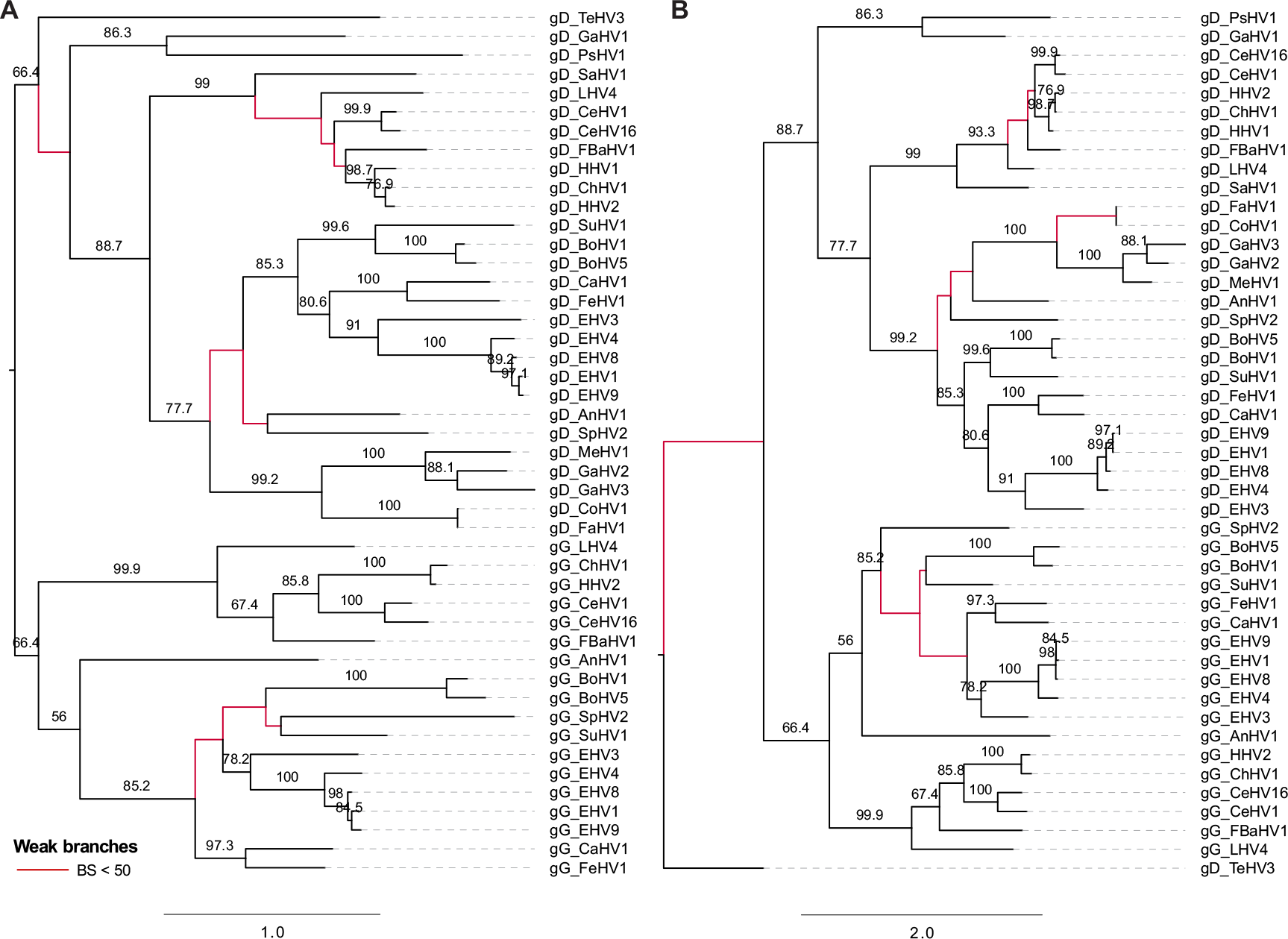
Glycoproteins D/G gene trees. A) Original tree inferred using PhyML, mid-rooted. B) Optimized gD/gG gene tree, where a new rooting and branching patterns were proposed after the first reconciliation step, as shown in Figure S1.

